# Novel kinetoplastid-specific cAMP binding proteins identified by RNAi screening for cAMP resistance in *T. brucei*

**DOI:** 10.1101/2023.03.14.532707

**Authors:** Sabine Bachmaier, Matthew K. Gould, Eleni Polatoglou, Radoslaw Omelianczyk, Ana E. Brennand, Maha A. Aloraini, Jane C. Munday, David Horn, Michael Boshart, Harry P. de Koning

## Abstract

Cyclic AMP signalling in trypanosomes differs from most eukaryotes due to absence of known cAMP effectors and cAMP independence of PKA. We have previously identified four genes from a genome-wide RNAi screen for resistance to the cAMP phosphodiesterase (PDE) inhibitor NPD-001. The genes were named cAMP Response Protein (CARP) 1 through 4. Here, we report an additional six CARP candidate genes from the original sample, after deep sequencing of the RNA interference target pool retrieved after NPD-001 selection (RIT-seq). The resistance phenotypes were confirmed by targeted RNAi knockdown and highest level of resistance to NPD-001, approximately 17-fold, was seen for knockdown of CARP7 (Tb927.7.4510). CARP1 and CARP11 contain predicted cyclic AMP binding domains and bind cAMP as evidenced by capture and competition on immobilised cAMP. CARP orthologues are strongly enriched in kinetoplastid species, and CARP3 and CARP11 are unique to *Trypanosoma*. Localization data and/or domain architecture of all CARPs predict association with the *T. brucei* flagellum. This suggests a crucial role of cAMP in flagellar function, in line with the cell division phenotype caused by high cAMP and the known role of the flagellum for cytokinesis. The CARP collection is a resource for discovery of unusual cAMP pathways and flagellar biology.

**Importance:** Trypanosomes are major pathogens of humans and livestock. In addition they have been invaluable as a model system to investigate new biological systems, and not just of protozoa. Equally, they are known to have a lot of unique biology and biochemistry. One example of this is signal transduction by cyclic nucleotides. Some elements, including phosphodiesterases and the catalytic domains of its dozens of adenylate cyclase isoforms, are highly conserved, while the absence of G-proteins, a cAMP-responsive protein kinase A and other known effector types suggests a unique cAMP-dependent pathway, which as yet is mostly uncharacterised. Here, we identify a set of ten *Trypanosoma brucei* proteins, all localised to its flagellum, that appear to be involved in the production of cAMP, or in mediating its cellular effects. These cAMP Response Proteins (CARPs) were mostly unique to trypanosomes, suggesting a completely novel pathway. Two of the CARPs were shown to bind cAMP and were found to possess structurally conserved cyclic nucleotide binding domains.

## Introduction

Conservation of signalling pathways and proteins among different phyla of eukaryotes is very limited, particularly in protozoa. Cyclic nucleotides are present as second messengers in almost all organisms, and in protozoa they can regulate growth, development and metabolic adaptation. In trypanosomes an important role of cAMP is suggested by an estimated 80 adenylate cyclase encoding genes (Salmon *et al*., 2012b), many of which are expressed throughout the life cycle (Alexandre *et al*., 1996); multiple are present in the plasma membrane (Bridges *et al*., 2008) or mainly localised to the flagellar surface (Paindavoine *et al*., 1992, Saada *et al*., 2014). These cyclases consist of a conserved catalytic domain, a single trans-membrane domain and a variable extracellular domain (Salmon, 2018). It is thus possible that various extracellular ligands control their activity (Paindavoine *et al*., 1992), but none have been identified and activation by dimerization has been observed under acidic, hypotonic or other stress conditions (Gould & de Koning, 2011, Nolan *et al*., 2000, Salmon, 2018). In recent years, the adenylate cyclases have been implicated in cytokinesis (Salmon *et al*., 2012a), immune evasion in the mammalian host (Salmon *et al*., 2012b), and social motility in the procyclic insect stage of the parasite (Lopez *et al*., 2015, Oberholzer *et al*., 2015, Saada *et al*., 2015). Control of cAMP homeostasis is crucial as knockdown of the locus containing the cAMP phosphodiesterases TbrPDEB1 and TbrPDEB2 is lethal (Oberholzer *et al*., 2007) and inhibitors of these enzymes have potent antitrypanosomal activity (de Heuvel *et al*., 2019, de Koning *et al*., 2012). However, the effectors and cascades activated or inhibited by cAMP remain largely unknown in trypanosomes, given the absence of genes encoding the known cAMP effectors in mammalian cells. Most importantly, the homologue of Protein Kinase A (PKA) is not activated by, and does not bind, cyclic nucleotides in *T. brucei* (Bachmaier & Boshart, 2014, Bachmaier *et al*., 2019, Bubis *et al*., 2018). This has stimulated a screen aiming at identification of novel cAMP effectors or target proteins. We previously reported a set of four *T. b. brucei* genes whose RNAi repression protected against the high intracellular cAMP concentration resulting from the inhibition of TbrPDB1/2; these genes were termed cAMP Response Protein (CARP) 1 – 4 (Gould *et al*., 2013). Out of these, only CARP3, a gene unique to trypanosomes, has been investigated and it was shown to be essential for social motility of procyclic trypanosomes through direct interaction with and regulation of adenylate cyclases (Bachmaier *et al*., 2022). CARP3 is therefore an upstream regulator of cAMP signalling and the effectors and cAMP-binding protein(s) remain to be identified.

The *Trypanosoma brucei* subspecies *T. b. gambiense* and *T. b. rhodesiense* are responsible for the disease known as Human African Trypanosomiasis (HAT), or sleeping sickness (Büscher *et al*., 2017), whereas *T. b. brucei* and the related trypanosomes *T. congolense*, *T. vivax*, *T. evansi* and *T. equiperdum* all contribute to the various manifestations of Animal African Trypanosomiasis (AAT), known under names such as nagana, surra and dourine (Giordani *et al*., 2016). Sleeping sickness has long been among the most neglected diseases, being transmitted by the tsetse fly, which makes it a problem of rural Africa, but more recently progress has been made and case numbers have dropped with more active control measures (Barrett, 2018, Franco *et al*., 2020) and the introduction of the first oral drug, fexinidazole, against the infection (Lindner *et al*., 2020). In contrast, little progress has been made in reducing the impact of AAT, which continues to have devastating effects on livestock and, consequently, on rural economies and food security in Africa and beyond, in part because *T. evansi*, *T. vivax* and *T. equiperdum* are not dependent on tsetse fly transmission (Aregawi *et al*., 2019, Desquesnes, 2004, Desquesnes *et al*., 2013, Jones & Dávila, 2001). No new animal trypanosomiasis drugs have been introduced for decades and the treatment options are both limited and threatened by resistance (Delespaux & de Koning, 2007, Giordani *et al*., 2016, Kasozi *et al*., 2022, Melaku & Birasa, 2013). Among the issues that have held back drug development against trypanosomiasis is the paucity of well-validated drug targets – a result of the many gaps in our knowledge of their unique biochemistry and cell biology. The limited understanding of mostly non-conserved signalling mechanisms and proteins is one reason why these pathways, preferred targets in mammalian drug development, are less exploited for the parasites.

Here we revisit the screen that identified CARP1 - 4 by deep sequencing of the RNAi library grown out after challenge with PDE inhibitor NPD-001 (Gould et al., 2013). We identify six additional genes that confer resistance to perturbation of intracellular cAMP, CARP6 – CARP11, and validate their phenotype by targeted RNAi for each individual gene. Most of the CARPs are unique to the kinetoplastidae. CARP1 and CARP11 contain potential cAMP-binding domains and cAMP binding to these proteins was confirmed by capture on cAMP-linked agarose beads. The identification of novel and trypanosomatid-specific cAMP effector candidates will elucidate the evolution of cAMP signalling and eventually provide attractive drug target candidates.

## Results

### 1. RNA-interference Target sequencing (RIT-seq) after selection for resistance to NPD-001

As described by (Gould *et al*., 2013), a genome-wide RNAi library screen was carried out with bloodstream *T. b. brucei* under selection pressure with the phosphodiesterase inhibitor NPD-001, previously known as CpdA (de Koning *et al*., 2012), following established protocols (Alsford *et al*., 2012, Alsford *et al*., 2011). In the initial report, we identified four genes by sequencing PCR-fragments from amplified prominent bands on an agarose gel and designated them CARP1 - 4. The sample from this selection has now been subjected to RIT-seq deep sequencing. This yielded a far more diverse and quantitative data set of genes that, upon RNAi knockdown, protect to any degree against the strongly elevated levels of intracellular cAMP resulting from the treatment with NPD-001. A total of 1183 genes was included in this dataset, with raw total counts between 74,563,689 and 1, out of a total of 105,662,532 bases mapped to CDS. Among the most frequent hits were CARP1 – CARP4 and an additional 7 genes were selected from the genes with highest counts, and designated CARP5 - CARP11 (Table 1). In addition, four adenylate cyclase-encoding genes were identified with high abundance, consistent with cyclase knockdown causing reduction of cAMP levels (Salmon *et al*., 2012a).

**Table 1.** Summary of cAMP response genes (CARPs) conferring cAMP sensitivity.

| Name | Tb427 accession<sup>1</sup> | Tb927 accession<sup>1</sup> | Length in<br>Tb927(a.a.) | % of mapped<br>RIT-seq bases | RNAi targeting<br>region (nt) | Localisation of mNeoGreen fusion<br>(tryptag.org; (Dean *et al.*, 2017)) |
| --- | --- | --- | --- | --- | --- | --- |
| CARP1 | Tb427_110181700 | Tb927.11.16210 | 705 | 70.6 | Gould et al., 2013 | Axoneme (cytoplasm, weak) |
| CARP2 | Tb427_110145400 | Tb927.11.12860 | 302 | 0.42 | Gould et al., 2013 | Cytoplasm (reticulated, weak)<sup>2</sup> |
| CARP3 | Tb427_070059100 | Tb927.7.5340 | 498 | 2.4 | Gould et al., 2013 | Flagellar tip, strong (flagellar pocket) |
| CARP4 | Tb427_030010200 | Tb927.3.1060 | 779 | 13.9 | Gould et al., 2013 | No information |
| CARP5 | Tb427_100020600 | Tb927.10.1740 | 382 | 3.38 | 365-776 | Nucleoplasm |
| CARP6 | Tb427_100131300 | Tb927.10.12390 | 258 | 4.7 | 256-682 | Axoneme |
| CARP7 | Tb427_070050500 | Tb927.7.4510 | 407 | 0.092 | 625-1177 | Axoneme (points in cytoplasm) |
| CARP8 | Tb427_110044600 | Tb927.11.3910 | 1108 | 1.1 | 2076-2624 | Axoneme (points in cytoplasm, weak) |
| CARP9 | Tb427_080051400 | Tb927.8.4640 | 385 | 0.25 | 601-1092 | Axoneme |
| CARP10 | Tb427_110076700 | Tb927.11.7180 | 653 | 0.34 | 2-439 | Axoneme (cytoplasm, weak) |
| CARP11 | Tb427_070027800 | Tb927.7.2320 | 605 | 0.046 | 524-990 | Axoneme, cytoplasm |

<sup>1</sup>Gene IDs are from TriTrypDB.org
<sup>2</sup>Flagellar localization or function likely from additional evidence discussed in the text

Figure 1A shows a pie-chart representation of the % of total mapped bases for each of the CARPs and adenylate cyclases that together account for 98.2% of all mapped bases in the RIT-seq analysis. Clearly, the most frequent gene targeted was CARP1, with 70.6% of all mapped bases, followed by CARP4 with 13.9%.

**Figure 1.**
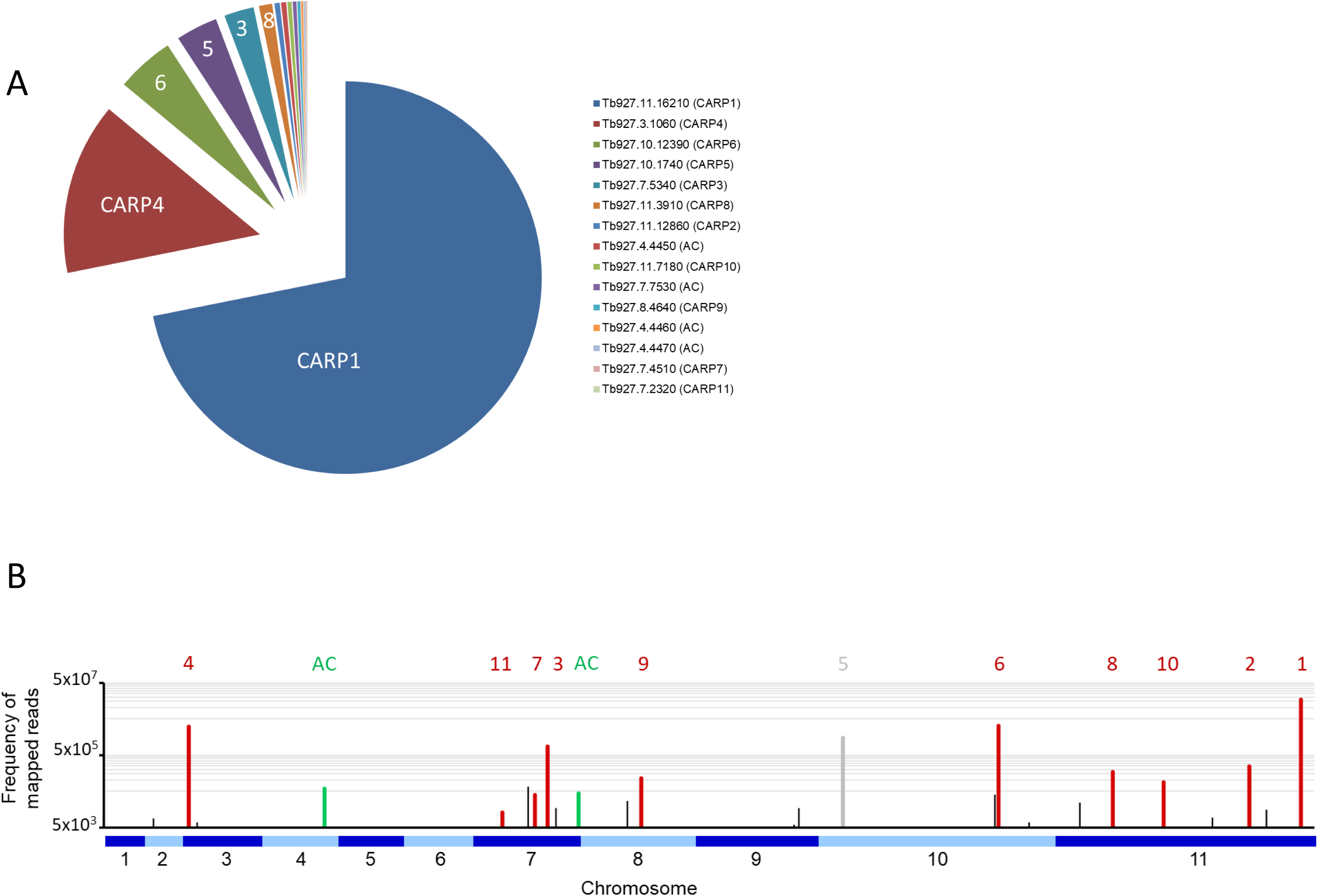
A. Pie chart of the percentage of all bases mapped to each of the eleven CARP genes and to the four GRESAG4 adenylate cyclases (AC). The gene identifiers from TriTrypDB and the CARP numbering are indicated. B. Frequency of mapped RIT-seq reads indicated on a genome map of T. brucei 927. Numbered CARPs are indicated by red bars and the corresponding numbers above them; only CARP5 is indicated in grey to indicate this was a false positive hit. Adenylate cyclases (AC) are similarly indicated in green; The green bar on chromosome 4 is an amalgamation of three bars, for genes Tb927.7.4450, Tb927.7.4460 and Tb927.7.4470. Black bars represent genes that were not followed up for validation by targeted RNAi for this study.

Of the newly identified CARPs, CARP6 was the most-targeted gene and CARP11 the least targeted, accounting for 4.69% and 0.046% of mapped bases, respectively. The four listed adenylate cyclases were identified between 0.42% and 0.14% of mapped bases. A genome map of RIT-seq hits is depicted in Figure 1B, which shows the designated CARPs, as well as the positions of the adenylate cyclases and other ‘hits’ that were not taken forward for verification by targeted RNAi. A listing of all 44 genes with normalised RIT-seq mapping ≥ 0.05 is given as Supplementary Table 1.

### 2. Phenotype confirmation of the new CARPs by targeted RNAi of the individual genes

Previously, we confirmed that targeted RNAi knockdown of *CARP1 − CARP4* did confer protection to treatment with NPD-001, i.e. a shift to a significantly higher EC_50_ in our standardised resazurin-based drug sensitivity test. This induced resistance was specific, as it was not afforded to treatment with control drugs pentamidine, suramin and difluoromethylornithine (Gould *et al*., 2013).

RNAi target sequences were chosen for each of the genes *CARP5 − CARP11*, in order to obtain true independent confirmation. The RNAi target fragments (Table 1) were cloned into the expression vector p2T7-177-BLE (Wickstead *et al*., 2002) and transfected into the *T. b. brucei* cell line Lister 427 MiTat 1.2 13-90 (Wirtz *et al*., 1999) for tetracycline (TET)-inducible expression. EC_50_ values were obtained in parallel with and without TET induction, allowing statistical comparisons. Further, the NPD-001 sensitivity tests were performed with four biological replicates (independent transfectants) and each EC_50_ was obtained on at least three separate occasions. Table 2 shows a summary of the CARP5 − CARP11 results, comparing the EC_50_ values obtained with and without TET induction. The RNAi knockdown resulted in significantly higher NPD-001 EC_50_ values for *CARP6* − *CARP11*, but not for *CARP5*, which may therefore be classified as a false positive from the library screen. Indeed, further analysis showed this gene to be a frequent false hit in RIT-seq owing to the RNAi vector junction sequence GCCTCGCGA. Mis-priming from sequence GGGCCAGT within the Tb927.10.1740 (*CARP5*) CDS produced a 1245-bp product. For the confirmed CARPs 6-11, the level of protection against elevated cAMP by targeted RNAi was by far highest for *CARP7* (17-fold), whereas the targeted knockdown of the other genes typically resulted in < 4-fold resistance gain (Table 2). The difference in response to knockdown for CARP5 and CARP7 is illustrated in Figure 2 with representative curves from the resazurin-based assays.

**Figure 2.**
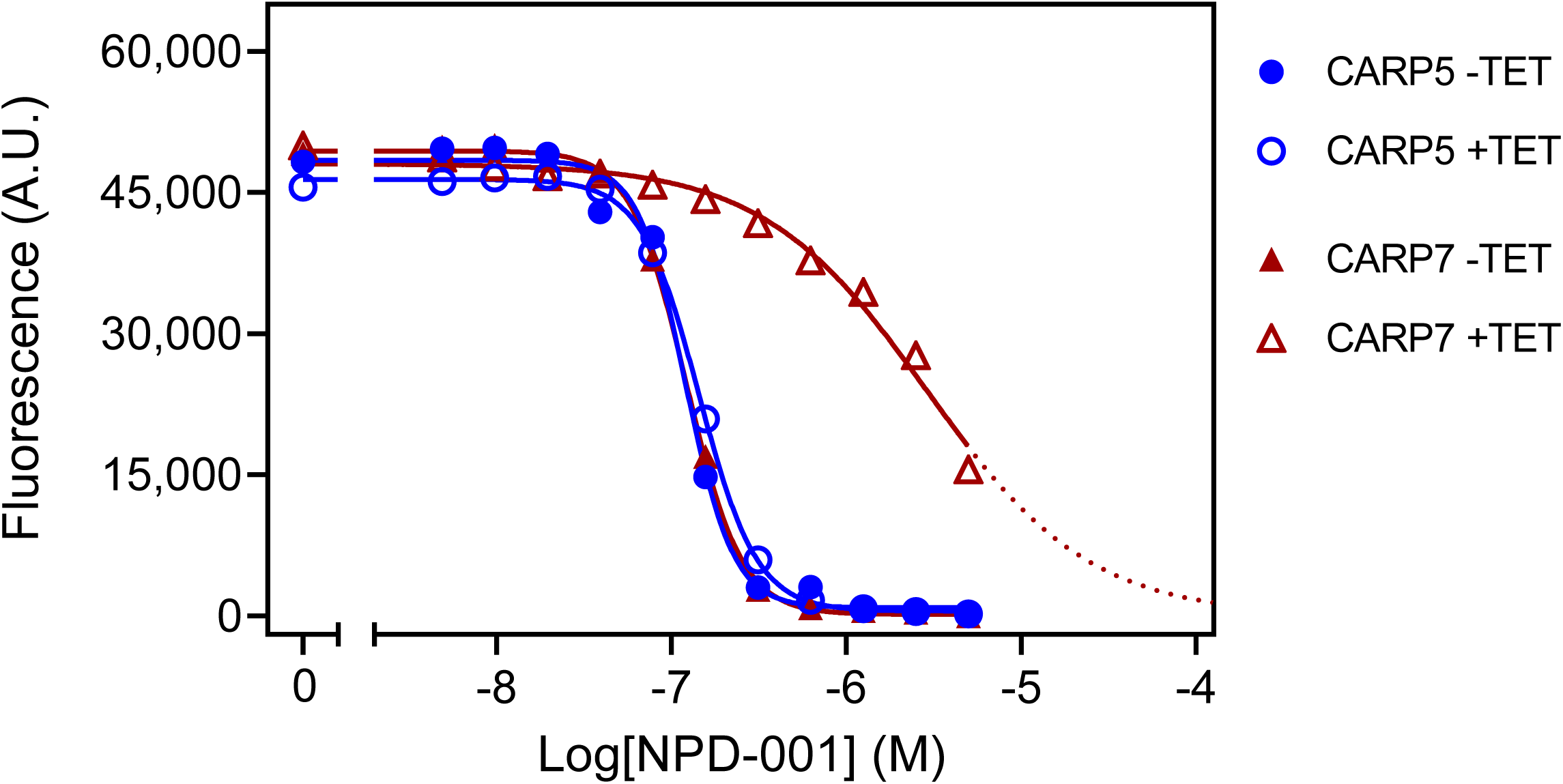
Representative graphs of the Alamar blue-based assay used to determine EC_50_ values for NPD-001. Dilution range for NPD-001 started at 5 μM and consisted of 11 doubling dilutions to 4.88 nM, and a no-drug control. The results shown are from a single experiment, representative of three independent replicates, performed with one of three clonal lines for each RNAi line (CARP5 clone 6 and CARP7 clone 4). Fluorescence read-outs in arbitrary units (A.U.), with background fluorescence subtracted, were plotted to a sigmoid curve with variable slope in Prism 9 to determine the EC_50_ values. Closed symbols represent the control without tetracycline (TET), open symbols, with TET-induction. For the TET-induced CARP7 values the dotted line indicates extrapolation.

**Table 2.** Sensitivity to phosphodiesterase inhibitor NPD-001 upon inducible RNAi

| RNAi target | EC<sub>50</sub> NPD-001<br>(Mean ± SEM), µM |  | Resistance<br>factor | *P*-value |
| --- | --- | --- | --- | --- |
|  | - Tet | + Tet |  |  |
| parental<br>control (n = 6) | 0.14 ± 0.01 | 0.17 ± 0.01 | 1.18 | 0.0369 |
| CARP5 (n = 9) | 0.14 ± 0.01 | 0.15 ± 0.01 | 1.09 | 0.0988 |
| CARP6 (n = 12) | 0.25 ± 0.04 | 0.61 ± 0.02 | 2.41 | <0.0001 |
| CARP7 (n = 12) | 0.22 ± 0.03 | 3.70 ± 0.27 | 16.91 | <0.0001 |
| CARP8 (n = 12) | 0.21 ± 0.01 | 0.41 ± 0.02 | 1.98 | <0.0001 |
| CARP9 (n = 12) | 0.26 ± 0.03 | 0.44 ± 0.08 | 1.72 | 0.0238 |
| CARP10 (n = 9) | 0.17 ± 0.01 | 0.30 ± 0.01 | 1.75 | <0.0001 |
| CARP11 (n = 12) | 0.18 ± 0.01 | 0.30 ± 0.01 | 1.69 | <0.0001 |

n = total number of replicates; *P*-value was calculated by unpaired, one-sided t-test.

### 3. Phylogeny and species distribution of CARP genes

CARP phylogeny was investigated by individual BLAST searches (blast.ncbi.nlm.nih.gov/Blast.cgi) as well as orthologue analysis based on the orthoMCL database (orthomcl.org) (Figure 3). Despite some small differences between the results of these two methods, all CARPs appear to be present in both African and American *Trypanosoma* species (*T. brucei*, *T. congolense*, *T. vivax*, *T. cruzi*) and most are also found in *Leishmania* genomes, except CARP3 and CARP11. CARP3 is also absent from the genomes of the insect gut parasites *Crithidia fasciculata* and *Angomonas deanei* as well as the free-living trypanosomatid *Bodo saltans*. CARP11 is found in *A. deanei* and *B. saltans* but no orthologue could be identified in *C. fasciculata*.

**Figure 3.**
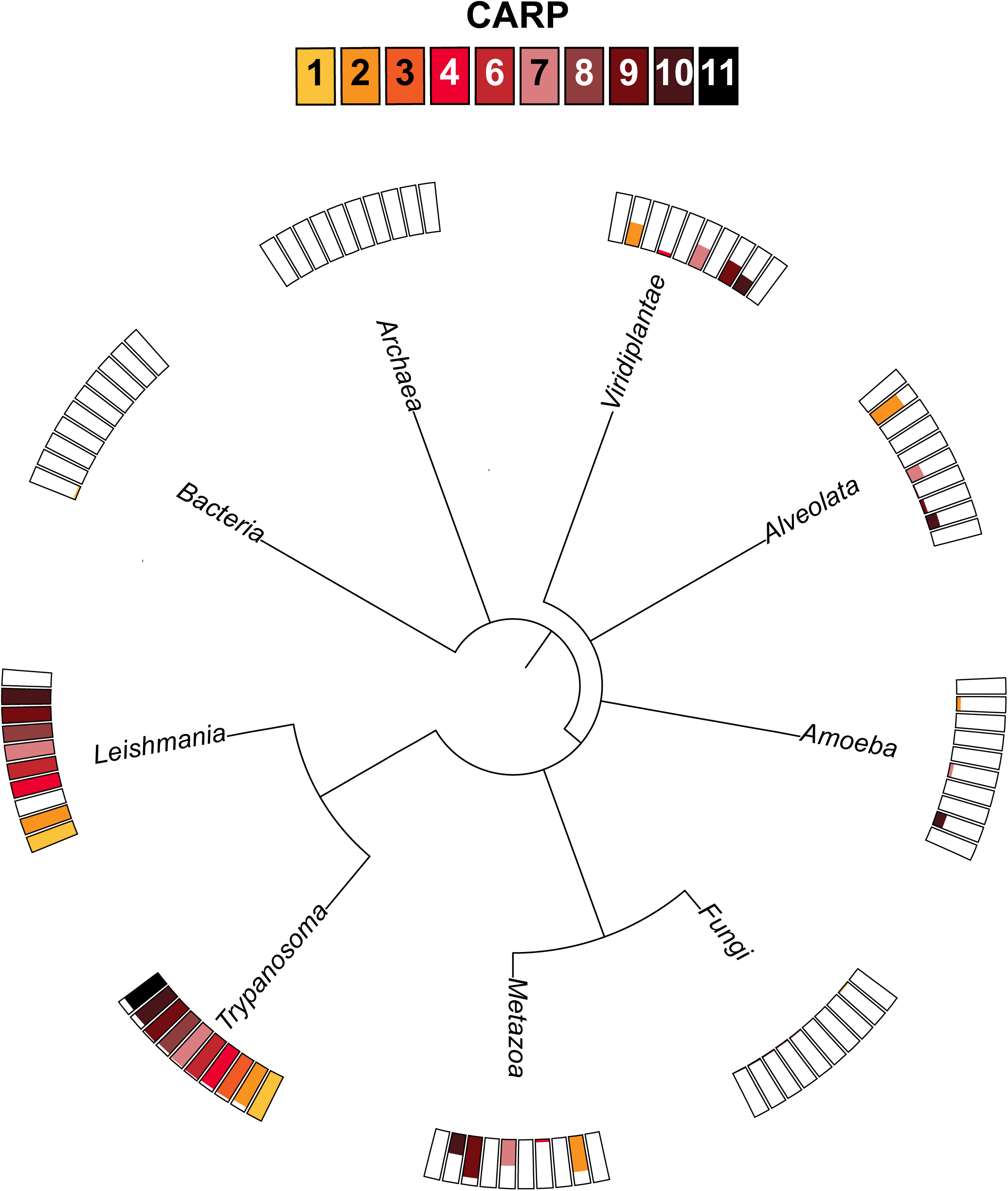
Distribution of ortholog groups for CARPs based on by OrthoMCL release 6.11 (https://orthomcl.org/orthomcl/app). Ortholog group IDs for CARPs were retrieved from TritrypDB release 58 (https://TritrypDB.org) and the tree was generated with phyloT version 2022.3 (https://phylot.biobyte.de/). Each CARP is shown by a differently coloured box and the size of the coloured bar represents the percentage of occurrence in the respective group of species.

Most CARPs are scarcely found outside the trypanosomatids, although there are some exceptions. CARP2 and CARP7 are found in some species of Viridiplantae, including the green algae *Chlamydomonas*, in some Alveolata, specifically the Apicomplexa, and in some metazoa including Mammalia; CARP9 is also present in *Chlamydomonas* and is quite widespread in metazoa, but is only found in very few Alveolata. Conversely, CARP4 is present in a few species of Viridiplantae and Metazoa but not in other non-trypanosomatids. Only CARP10 had a few orthologues in Amoebozoa and Alveolata and some in Viridiplantae and Metazoa. None of the CARPs had detectable orthologues in prokaryotes, fungi or in *Arabidopsis*. Overall, most CARPs are (almost) exclusive to trypanosomatids, with CARP3 limited to the genus *Trypanosoma*.

### 4. Analysis of domain structure and subcellular localisation of the CARP proteins

The domain structure and subcellular localization of the 10 confirmed CARPs (i.e., excluding CARP5) is summarised in Figure 4. Domains could be identified in profile database searches for all except CARP8, although some of these are Domains of Unknown Function (DUF), specifically DUF4464 as the main conserved domain in CARP2 and DUF4201 as the only domain detected in CARP7.

**Figure 4.**
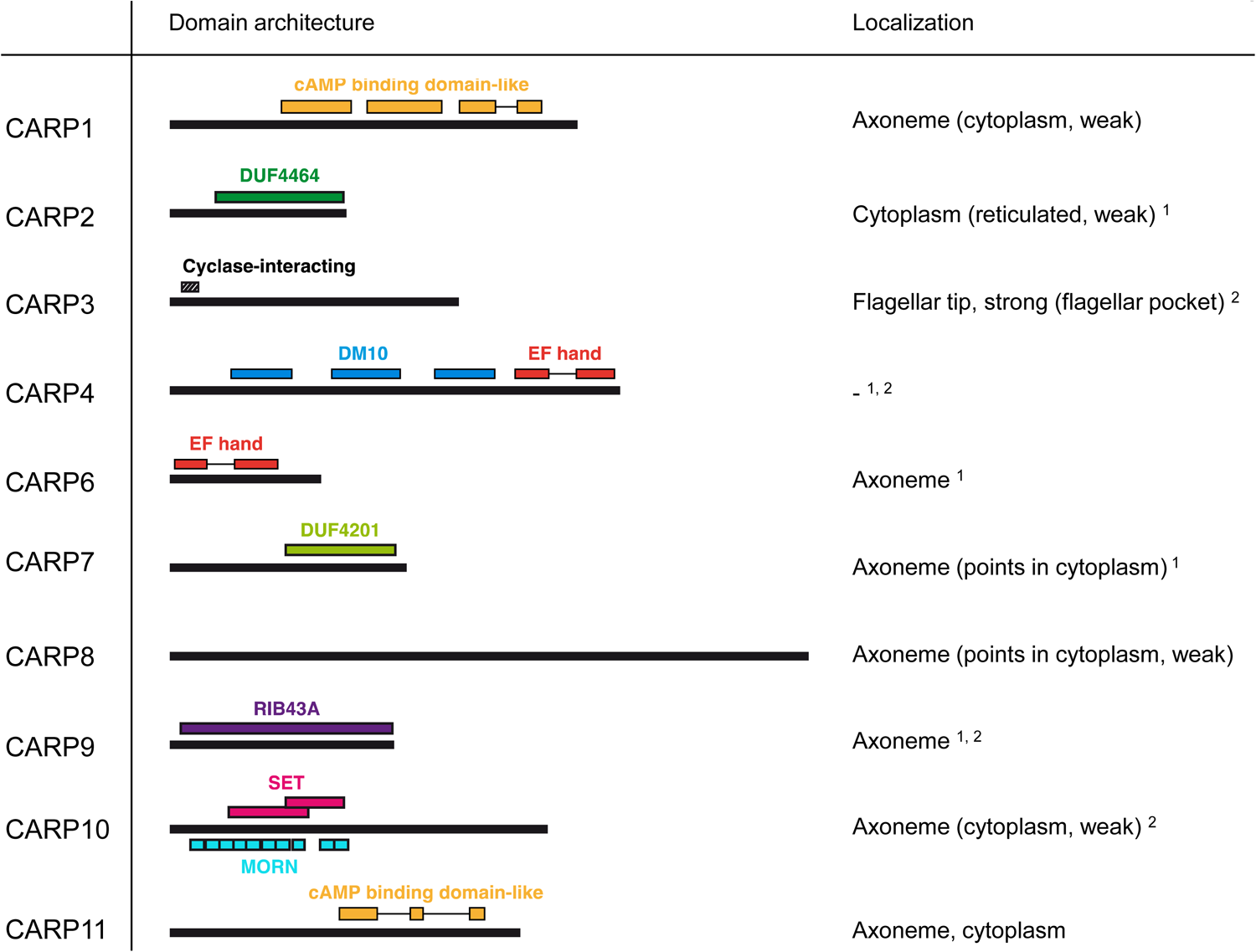
Domain predictions for CARPs by the domain profile databases Superfamily 2.0 (https://supfam.org/), Pfam version 35.0 (http://pfam.xfam.org/), and/or SMART (http://smart.embl-heidelberg.de/) are shown as schematic representation. CARP proteins are shown as black bars (to scale) and predicted domains are displayed by coloured boxes with IDs as indicated in the figure. For CARP3, the region required for flagellar tip localization and cyclase interaction (designated ‘Cyclase-interacting’) according to Bachmaier *et al*. (2022) is indicated. Subcellular localization of CARP proteins is given exactly as annotated in the TrypTag.org database (Dean *et al*., 2017). Additional evidence for axonemal^1^ or flagellar^2^ localization is taken from Broadhead *et al*. (2006) and Subota *et al*. (2014), respectively.

Strikingly, the TrypTag genome-wide tagging project (Billington *et al*., 2023, Dean *et al*., 2017) revealed an axonemal localization for 8/9 CARPs successfully tagged, which closely aligned with Gene Ontology (GO) assignments in TriTrypDB (Supplementary Table S1). The exception was CARP2, for which no GO assignments were available, and which showed a cytoplasmic distribution as N-terminal mNeonGreen fusion protein in TrypTag. However, CARP2 and CARPs 3-7, 9 and 10 are all present in at least one flagellar proteome of procyclic trypanosomes (Broadhead *et al*., 2006, Subota *et al*., 2014). Also, many CARP2 orthologues from other organisms are annotated as cilia- and flagella-associated proteins, and a bioinformatic study by Baron *et al*. (2007), identified CARP2 as a conserved component of motile cilia and flagella, supporting a flagellar localization. The same study also identified CARP4, the only CARP without localization information on TrypTag.org (accessed 02/04/2023). CARP4 is orthologous to *Chlamydomonas reinhardtii* Rib72 (Chlre5_6|10141; BLASTP E-value = 3.1E-28), which appears to be involved in linking microtubules of the outer doublets of the axoneme (Ikeda *et al*., 2003). Like CARP4, CrRIB72 contains three DM10 domains and a C-terminal EF-hand domain, which binds Ca^2+^ and is implicated in regulatory and signalling responses (Nelson *et al*., 2002). Moreover, CARP4 is orthologous to the human EF Hand Containing protein EFHC1 (BLASTP E-value = 5E-55) that is linked to Juvenile Myoclonic epilepsy (King, 2006) and is believed to be part of the structure of cilia (King, 2006, Suzuki *et al*., 2020). CARP9 is annotated as Component of Motile Flagella 19 (CMF19) in TriTrypDB. It contains a RIB43A domain, a structure that was first identified in *Chlamydomonas*, and is associated with protofilament ribbons and basal bodies of ciliary and flagellar microtubules (Norrander *et al*., 2000). As the microtubular ribbons in *Chlamydomonas* mostly consist of RIB43A and RIB72 in addition to tubulin (King, 2006), it is likely that CARP4 and CARP9 play a similar role in *T. brucei*, with the EF-hand domain providing Ca^2+^-mediated regulation. Curiously, CARP9 was co-purified with the *T. brucei* Mitochondrial Calcium Uniporter (TbMCU; (Huang & Docampo, 2020)), although it is not immediately clear how a putative axonemal protein and an integral membrane protein of the inner mitochondrial membrane might associate *in vivo*. CARP6 also contains an EF-hand domain (Figure 4), reinforcing the link to Ca^2+^ signalling somewhere in the response to high cAMP.

CARP3’s predicted structure shows a highly structured N-terminus essential for localization of the protein at the distal tip of the flagellum in procyclic trypanosomes (Bachmaier *et al*., 2022) where it interacts with several adenylate cyclases and regulates social motility and tsetse fly transmission (Bachmaier *et al*., 2022, Shaw *et al*., 2022).

CARP10 contains a series of ten predicted Membrane Occupation and Recognition Nexus (MORN) domains as well as two copies of the Histone H3 K4-specific methyltransferase SET7/9 N-terminal domain superfamily (101-220 and 201-300). Although the MORN domain has been identified in a total of 113,928 proteins in many species (SMART database; http://smart.embl.de/smart/; 25/07/2022), the function of this domain is still poorly understood (Zhou *et al*., 2022). It localises to the axoneme based on GO terms and TrypTag.

Finally, cAMP binding motifs were identified in the sequences of two CARP proteins, CARP1 and CARP11, i.e. the genes to which the highest and the lowest percentage of reads, respectively, were mapped in the RIT-seq screen. In CARP1, three cAMP binding domains are detected at a.a. positions 233–346, 370–491 and 519–577,612–651, whereas in CARP11 a single cyclic nucleotide binding domain is detected, with insertions, at positions 293–358, 415–437 and 519–545.

The putative phosphate binding cassette (PBC) sequences of the predicted CARP1 and CARP11 cAMP binding domains were aligned with well-characterised but diverse cAMP binding domains from different species: *E. coli* CRP (EcCRP, Uniprot P0ACJ8), *Mus musculus* EPAC4 (MmEpac4, Uniprot Q9EQZ6) and *Bos taurus* PKARIα (BtPKARIα, Uniprot P00514) in Figure 5. A PBC consensus sequence that was compiled for PKA regulatory subunits, F-G-E-[LIV]-A-L-[LIMV]-x(3)-[PV]-R-[ANQV]-A (Canaves & Taylor, 2002, Kannan *et al*., 2007), is also displayed for comparison. Highly conserved residues of the PBC include an invariant arginine (R211 in BtPKARIα) known to interact with the phosphate group of cAMP and a glutamate (E202 in BtPKARIα) that interacts with the 2’OH group of the ribose (Berman *et al*., 2005). While CNBD-A of CARP1 and CARP11 match very well with the PKA consensus, CNBD-B and -C of CARP1 have important deviations and the validity of the alignment and cAMP binding prediction are questionable (Figure 5A, C). Structural alignment of the CNBDs in the AlphaFold2-predicted CARP1 structure (retrieved from http://wheelerlab.net/alphafold/) with published crystal structures of diverse cAMP binding proteins (EcCRP, PDB 4N9H; BtPKARIα, PDB 1RGS) showed a very good structural preservation including overlay of R320 in CARP1 CNBD-A with the conserved arginines of the reference protein structures (Figure 5B, RMSD 1.168 with EcCRP CNBD; RMSD 0.782 with BtPKARIα CNBD-A). The same observation is made for R520 in the CARP11 CNBD (Figure 5D, RMSD 1.039 with EcCRP CNBD; RMSD 0.602 with BtPKARIα CNBD-A). In contrast, the conserved arginine in the reference structures aligned with Lys459 and Tyr629 in CNBD-B and CNBD-C of CARP1, respectively (Supplemental Figure S1A, B).

**Figure 5.**
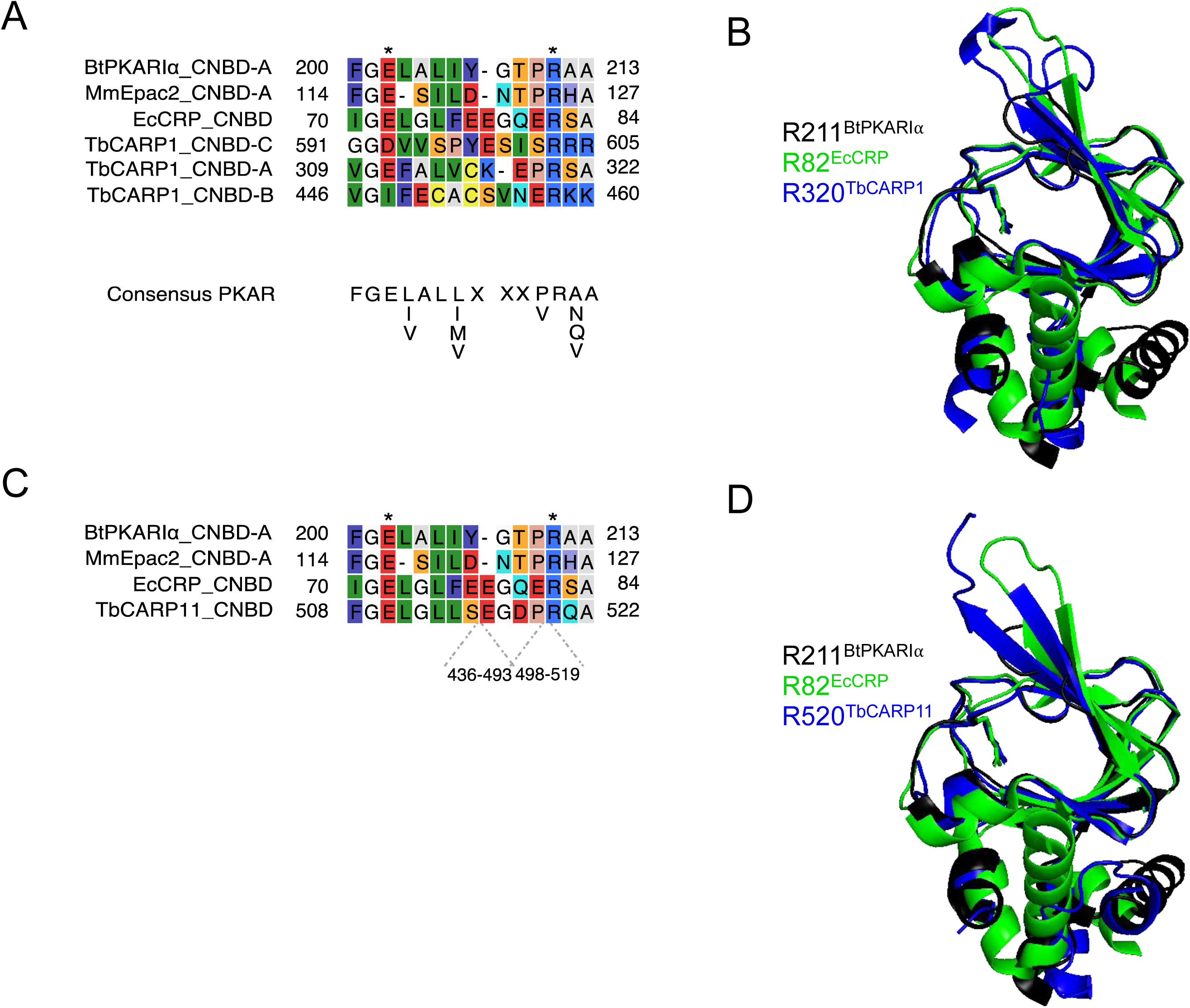
Sequence comparisons (A, C) of the predicted phosphate binding cassettes (PBCs) of *T. brucei* CARP1 (A, CNBD-A, -B, -C) or CARP11 (C) with *Escherichia coli* CRP (EcCRP), *Mus musculus* EPAC2 (MmEpac2) and *Bos taurus* PKARIα (BtPKARIα). The amino acid numbers of the predicted sequence insertions within the CARP11 PBC are indicated below the alignment. Asterisks indicate conserved Glu and Arg residues known to be essential for cAMP binding (Berman *et al*., 2005). (B, D) Structural alignments of CARP1 CNBD-A (B, dark blue) or CARP11 CNBD (D, dark blue) with BtPKARI*α* CNBD-A (PDB 1RGS, black) and EcCRP CNBD (PDB 4N9H, green) with the conserved arginines R320 of CARP1 and R520 of CARP11, respectively, overlaying the conserved arginines R211 of BtPKARIa CNBD-A and R82 of EcCRP CNBD. The structures were retrieved from http://wheelerlab.net/alphafold/ and superimposed with Pymol version 2.5.4.

### 5. Experimental validation of cAMP binding of CARP1 and CARP11

In order to test whether CARP1 and CARP11 do bind cAMP, we performed a pull-down experiment with cAMP immobilised on agarose beads via a flexible aminohexyl linker on position 8 (8-(6-Aminohexylamino)-adenosine-3’, 5’-cyclic monophosphate; 8-AHA-cAMP-agarose). Lysates of *T. brucei* overexpressing CARP1 were first incubated with agarose beads without attached cAMP in order to remove proteins non-specifically binding to the bead matrix. Subsequent incubation with 8-AHA-cAMP-agarose allowed the pulldown of a fraction that was highly enriched in cAMP-binding proteins. This procedure was performed in the presence of increasing cAMP concentrations (0 – 50 µM) intended to outcompete the binding to 8-AHA-cAMP. Subsequent elution of the protein fraction from the agarose beads, followed by western blot with a rabbit anti-CARP1 serum gave a quantifiable amount of CARP1 that was plotted against the cAMP concentration to derive an approximation of cAMP binding affinity. The antiserum had been validated using a panel of *T. brucei* cell lines, showing an increased signal in the CARP1-overexpressing strain and a decrease upon RNAi knockdown (Supplemental Figure S2). Figure 6A shows the average of two such determinations plotted with a sigmoidal curve with variable slope (standard inhibition; r^2^ = 0.977) yielding an EC_50_ value of 32.4 nM (95% CI 14.6 – 76.1 nM). A further pull-down with cAMP-agarose beads was performed using a set of differently linked cAMP analogues. Beads with various linkers connected to the purine ring at positions 2, 6, or 8 all showed similar capacity to pull down CARP1, but two different linkers attached to the 2’ position of the ribose failed to pull down any CARP1 (Supplemental Figure S3), consistent with cAMP interaction within the CNB pocket primarily via the ribose and cyclic phosphate.

**Figure 6.**
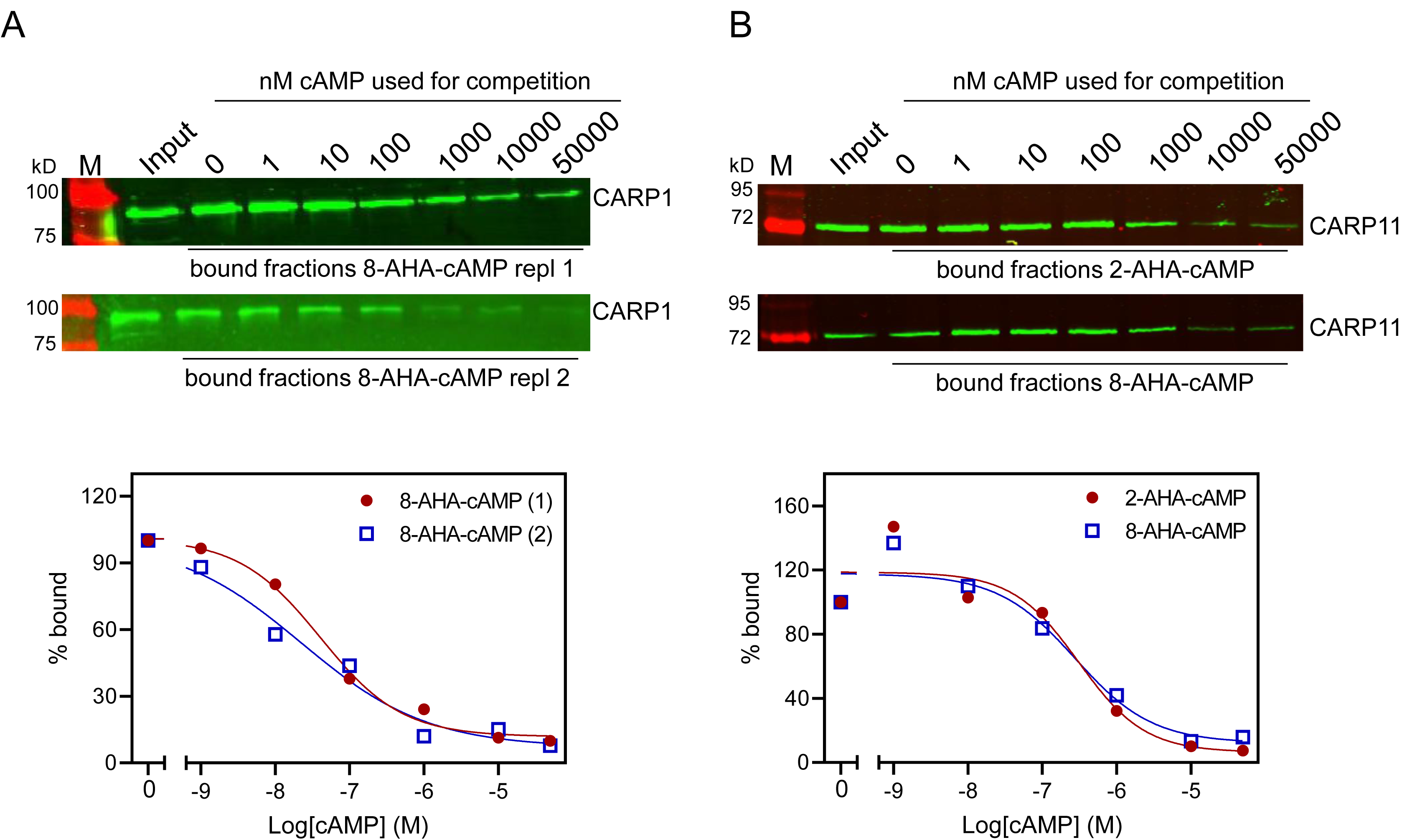
Cyclic AMP affinity purification of CARPs with predicted cAMP binding domains. (A) CARP1 was pulled down by 8-AHA-cAMP agarose from *T. brucei* cells overexpressing CARP1 (2 replicates). (B) CARP11 was expressed in *E. coli* with a C-terminal SUMO-His tag and pulled down by 2-AHA-(top) or 8-AHA-cAMP agarose (bottom). The western blots show the input fractions and the pulled down material in presence of increasing concentrations of cAMP (0 nM – 50 µM) for competition. Binding curves from the fluorescence quantifications are shown below; the amount of CARP pulled down without competition was set to 100%.

The same procedure was performed with CARP11 expressed with a SUMO-His tag in *E. coli* (Figure 6B), using 8-AHA-cAMP and 2-AHA-cAMP agarose beads. For western blots a mouse anti-His serum was used for detection of the tagged CARP11. The competition titrations with cAMP in Figure 6B show similar binding to 8-AHA-cAMP and 2-AHA-cAMP agarose beads with almost identical EC_50_ estimates: 309 nM for 2-AHA-cAMP and 276 nM for 8-AHA-cAMP. We conclude CARP1 and CARP11 are cAMP-binding proteins and that CARP1 has a ∼10x higher affinity for the cyclic nucleotide than CARP11.

## Discussion

By genome-wide RNAi screening for cAMP resistance in bloodstream stage trypanosomes, we have identified a set of proteins with a predicted function in cAMP metabolism or cAMP signalling. The selection was based on the essentiality of cAMP metabolism (Oberholzer *et al*., 2007, Salmon *et al*., 2012a, Shakur *et al*., 2011) and a strong cell division phenotype resulting from genetic or pharmacological perturbations of cellular cAMP concentrations (de Koning *et al*., 2012, Gould *et al*., 2013, Salmon *et al*., 2012a). Strikingly, for all identified cAMP response proteins (CARPs) we found publicly available experimental evidence (summarized in Fig. 4) for association with the flagellar axoneme or the flagellum, suggesting an important impact of cAMP signalling on flagellar biology. An intact and functional flagellum was previously shown to be essential for cell division of bloodstream stage trypanosomes, as RNAi repression of many genes with flagellar localization or function result in a lethal cell division phenotype (Broadhead *et al*., 2006; Marquez et al., 2022). This may explain why the screen returned specifically the observed set of proteins. Initiation of flagellar replication is the necessary first step in trypanosome cell division (Sun *et al*., 2013, Vaughan, 2010), and impaired flagellar function results in a block in cytokinesis. This is exactly the lethal phenotype seen with non-physiologically elevated cAMP (de Koning *et al*., 2012, Gould *et al*., 2013). Consequently, cyclic nucleotide phosphodiesterases (PDEs) of trypanosomes are considered good drug targets (Blaazer *et al*., 2018, de Heuvel *et al*., 2019).

A priori, we expected to get from our RNAi screening the following protein categories (1) proteins involved in cAMP production like adenylate cyclases (ACs) or proteins affecting activity or abundance of ACs, (2) effectors binding cAMP and (3) a diverse group of targets downstream of the effectors or modulating the activity of those effectors. As expected, the hit list included several AC genes, but at relatively low RIT-seq representation. The *T. brucei* genome contains more than 70 receptor-type ACs, the catalytic domains of which share homology with mammalian class III cyclases (Durante *et al*., 2020, Pays & Nolan, 1998, Salmon, 2018, Salmon *et al*., 2012b). Many of the *T. brucei* ACs have been shown to be developmentally regulated (Naguleswaran *et al*., 2018, Saada *et al*., 2014), while others are expressed throughout the life cycle (Alexandre *et al*., 1996). The fact that we observed only relatively few ACs, and at relatively low RIT-seq representation (Fig. 1), may be explained by the limited impact on the cAMP concentration upon RNAi of any individual AC family members (Salmon *et al*., 2012a). A higher RIT-seq representation was found for CARP3, a membrane-associated protein that interacts with several ACs, including the ESAG4 cyclase that is abundant in bloodstream forms, and positively regulates their abundance (Bachmaier *et al*., 2022). It is therefore likely that CARP3 repression confers cAMP resistance in bloodstream forms by concomitant downregulation of a significant fraction of ACs, particularly the dominant ESAG4, and thereby lower cAMP production.

At the effector level, we have identified two novel cAMP-binding proteins from the RNAi screen, CARP1 (Gould *et al*., 2013) and CARP11. The *T. cruzi* CARP1 homolog (TcCLB.508523.80) also seems to bind cAMP (Jäger *et al*., 2014). CARP1 has three predicted CNB domains; the phosphate binding cassettes (PBCs) of the first (CNBD-A) and of the CNB domain of CARP11 match a consensus sequence for the PBCs of PKA of diverse species very well (Fig. 5A, C). In contrast, CNBD-B and CNBD-C of CARP1 deviate in critical residues and therefore, we can neither predict nor exclude their contribution to cAMP binding in our immobilised cAMP assay (see also Fig. S1). Interestingly, CARP11 has two longer sequence insertions within the CNBD that apparently do not compromise the cAMP binding function. AlphaFold2 modelling of the domain and overlay with reference CNB crystal structures (Fig. 5D) show indeed a well conserved CNB structure in CARP11 and suggests that the insertions have accessory functions compatible with cAMP binding. The binding affinity, estimated by EC50 of cAMP competition, is ∼30 nM for CARP1, a value in the range of the free regulatory subunit of PKA from other organisms (Moll *et al*., 2007). The EC50 for CARP11 is ∼300 nM and closer to effectors like mammalian EPAC (Dao *et al*., 2006) and to physiologically relevant cAMP concentrations in cAMP microdomains (Anton *et al*., 2022).

The remaining seven CARP genes identified in the genome-wide RNAi screen cannot be assigned yet to a specific biochemical function. Some of them are most likely targets downstream of the cAMP signal, supported by the identified domains and available data on potential orthologs in higher eukaryotes (e.g. CARPs 2, 4, 9). For others (CARPs 6, 7, 8, 10), a function at the level of cAMP signal production, as found for CARP3, is equally possible. EF-hand domains (Nelson *et al*., 2002) in CARP4 and 6 suggest crosstalk between cAMP signalling and regulatory processes involving Ca^2+^.

Most importantly, the experimentally determined subcellular localization in trypanosomes retrieved from the database provided by the TrypTag project (Billington et al., 2023, Dean *et al*., 2017) assigned 7/10 CARPs to the flagellar axoneme and 2/10 (CARPs 2 and 4) were assigned to the flagellum based on proteome data (Broadhead *et al*., 2006, Subota *et al*., 2014). CARP3 is associated with the flagellar membrane and the axonemal cap via FLAM8 (Bachmaier *et al*., 2022). Thus, all CARPs are predicted to have a role in flagellar biology. By inference, cAMP signalling is important for regulation of flagellar functions. The impact is not limited to trypanosomes: 6/8 of the CARPs present in the *Leishmania* genome (Fig. 3) have been found enriched in the flagellar insoluble fraction by proteome analysis (CARPs 2, 4-6, 9, 10, (Beneke *et al*., 2019)). The CARPs thus provide a valuable resource to functionally dissect important flagellar biology of kinetoplastid parasites. Indeed the longer list of 44 genes with normalised RIT-seq mapped reads ≥ 0.05 (Supplementary Table S1) was highly enriched with axonemal (16/44) and other flagellar (7/44) localisations (TrypTag.org) and was significantly enriched for the GO component terms ‘axoneme’ (6-fold; P = 2.7×10^-8^), ‘ciliary plasm’ (2.3-fold; P = 7.7×10^-5^) and ‘cilium’ (1.85-fold; P = 5.7×10^-4^). This dataset could thus provide a starting point for further investigation into flagellar biology as well as novel cAMP-regulated pathways essential for parasite growth, survival and transmission. These pathways are potential targets for drug development against kinetoplastid diseases.

One example of an in-depth analysis is provided by CARP3. This protein localizes to the flagellar membrane in bloodstream forms but is restricted to a specialised microdomain at the tip of the flagellum in the procyclic stage (Bachmaier *et al*., 2022). In that stage, cAMP signalling via tip-localized adenylate cyclases (ACs) has been shown to regulate social motility “SoMo” on agarose plates (Lopez *et al*., 2015, Oberholzer *et al*., 2015), a phenotype that is now interpreted as chemotaxis along a pH gradient towards a more basic environment (Shaw *et al*., 2022). CARP3 interacts with multiple ACs in a flagellar tip complex and is essential for “SoMo” and for colonization of tsetse fly tissues and thus transmission (Bachmaier *et al*., 2022, Shaw *et al*., 2022, Shaw & Roditi, 2023). CARP3 differentially regulates the abundance of ACs, and a model predicts that it controls the relative abundance of functionally different cyclases for integration of various external signals (Bachmaier *et al*., 2022). The flagellar tip and membrane localization of CARP3 was also seen in *T. cruzi* (Won *et al*., 2023). The specific role in signalling from the environment to the cell, e.g. pH chemotaxis, is consistent with the membrane-associated localization of CARP3. In contrast, the other CARPs are localized to the axoneme and therefore more likely play roles in cell autonomous pathways, regulating flagellar motility. The fact that flagellar motility *per se* is unaffected in CARP3-depleted SoMo-negative cells (Bachmaier *et al*., 2022) also argues for distinct pathways. In mammalian cells, cAMP is involved in regulating ciliar motility within the axoneme (Wirschell *et al*., 2011) as well as distinct cilial sensing pathways (Nachury & Mick, 2019).

The novel cAMP-binding protein CARP1 appears in the genome of *Euglenozoa* (Fig. 3), also supported by detection of CARP1, 4, 6, 8, 9, 10 in the flagellar proteome of *Euglena gracilis* (Hammond *et al*., 2021). In contrast, CARP11 is only detected in *Trypanosoma*. We notice a striking correlation between the exclusive presence of these cAMP effectors in *Euglenozoa* and the absence of a cAMP-dependent protein kinase A. PKA in *T. brucei* and *T. cruzi* is structurally conserved but repurposed for binding and regulation by nucleoside analogs (Bachmaier *et al*., 2019). Future dissection of the novel flagellar cAMP signalling mechanisms may indicate why repurposing of PKA for a different second messenger or activation mechanism might have conferred a selective evolutionary advantage to organisms with novel cAMP effectors.

## Methods

### Trypanosome culture conditions

Bloodstream forms of the monomorphic *Trypanosoma brucei brucei* strain Lister 427 MiTat 1.2 were cultivated at 37 °C and 5% CO_2_ in modified HMI-9 medium (Vassella *et al*., 1997) supplemented with 10% (v/v) heat-inactivated foetal bovine serum (FBS). For maintenance of tetracycline repressor Tet^R^ and T7 polymerase in the trypanosome cell lines MiTat 1.2 13-90 or single marker (Wirtz *et al*., 1999), 2.5 µg/mL G418 and 5 µg/mL hygromycin B or only 2.5 µg/mL G418, respectively, were added to the culture medium. Cell density was monitored using a haemocytometer and was kept below 1 × 10^6^ cells/mL for continuous growth of replicative long slender bloodstream forms.

### RNAi target sequencing (RIT-seq)

Sequencing of the libraries was performed on the Thermo Scientific Ion Proton at the Glasgow Polyomics facility using the 200 base pair sequencing kit. sequences containing a terminal RNAi-vector junction sequence (GCCTCGCGA) were mapped to the *T. brucei* 927 reference genome (Alsford *et al*., 2012), the selection allowed for a single aberration, either a base change or a single base insertion or deletion. The filtered reads were mapped to the reference genome with Bowtie2 using the local mode alignment. The aligned reads were then assigned to a region of interest, reads that either were fully contained within the region or only partially overlap the region of interest. The counts for each replicate were expressed as normalised mapped reads by dividing by the CDS length × 100, in addition to percentage of total reads.

### Cloning and generation of transgenic trypanosomes

#### Generation of inducible CARP knock down cell lines

For tetracycline-inducible knock down of putative CARPs, the following fragments were amplified from *T. brucei* Lister 427 MiTat 1.2 single marker genomic DNA and cloned into p2t7-177-BLE (Wickstead *et al*., 2002) via BamHI and XhoI restriction sites: *CARP5* (Tb927.10.1740): RNAi targeting fragment: ORF nt 365-776, amplified with primers Tb427.10.1740_RNAi_Frag-F and Tb427.10.1740_RNAi_Frag-R; *CARP6* (Tb927.10.12390): RNAi targeting fragment: ORF nt 256-682, amplified with primers Tb427.10.12390_RNAi_Frag-F and Tb427.10.12390_RNAi_Frag-R; *CARP7* (Tb927.7.4510): RNAi targeting fragment: ORF nt 625-1177, amplified with primers Tb427.07.4510_RNAi_Frag-F and Tb427.07.4510_RNAi_Frag-R; *CARP8* (Tb927.11.3910): RNAi targeting fragment: ORF nt 2076-2624, amplified with primers Tb427tmp.02.1410_RNAi_Frag-F and Tb427tmp.02.1410_RNAi_Frag-R; *CARP9* (Tb927.8.4640, alternative name: CMF19, component of motile flagella 19): RNAi targeting fragment: ORF nt 601-1092, amplified with primers Tb427.08.4640_RNAi_Frag-F and Tb427.08.4640_RNAi_Frag-R; *CARP10* (Tb927.11.7180): RNAi targeting fragment: ORF nt 2-439, amplified with primers Tb427tmp.02.5030_RNAi_Frag-F and Tb427tmp.02.5030_RNAi_Frag-R; *CARP11* (Tb927.7.2320): RNAi targeting fragment: ORF nt 524-990, amplified with primers Tb427.07.2320_RNAi_Frag-F and Tb427.07.2320_RNAi_Frag-R. The RNAi plasmids were linearized with NotI for transfection and cells were selected by addition of 2.5 µg/mL phleomycin to the culture medium.

#### Inducible overexpression of CARP1

The CARP1 ORF was amplified by PCR from *T. brucei* MiTat 1.2 genomic DNA using primers Tb11.01.7890 SfiI up and Tb11.01.7890 BamHI low and cloned into plew82 (Wirtz *et al*., 1999) via SfiI and BamHI. Point mutations in the predicted CARP1 cyclic nucleotide binding domains were generated by site-directed mutagenesis using primers Tb11.01.7890 CNBDmut1 up and Tb11.01.7890 CNBDmut1 low (R320L, AGA → TTA), Tb11.01.7890 CNBDmut2 up and Tb11.01.7890 CNBDmut2 low (R458L, CGA → CTA), or Tb11.01.7890 CNBDmut3_R605L up and Tb11.01.7890 CNBDmut3_R605L low (R605L, AGG → CTG), respectively. The plasmids were linearized with NotI for transfection and cells were grown in presence of 2 µg/ml blasticidin.

#### Recombinant protein expression in *E. coli*

The CARP11 ORF (Tb927.7.2320) was amplified from *T. brucei* MiTat 1.2 genomic DNA with primers Tb927.7.2320_SUMO_BamHI_fw (GCACTAGGATCCATGAGCATCGTCAATCAG) and Tb927.7.2320_SUMO_HindIII_rev (CCACCAAGCTTTCACTTGACACGACTGCA) and cloned into pETM11_SUMO3 via BamHI and HindIII. The protein was expressed in *E. coli* Rosetta (grown in TB with 1 M sorbitol, overnight induction of protein expression with 200 µM IPTG at 16 °C) and the soluble fraction was used for pull-down with cAMP agarose.

### Generation of polyclonal antibodies

The *CARP1* ORF was amplified using primers Tb427tmp.01.7890.fl.f.10His (ataccatgggccaccaccaccaccaccaccaccaccaccacggcgcgggcGGTAGTTATGAATACCCAGACTAC) and Tb427tmp.01.7890.fl.b (aattcggatcctggctTTACCTTTTCGCCATGAACTCTTT) introducing an N-terminal His_10_ tag and cloned into pETDuet-1 via NcoI and BamHI restriction sites. The protein was expressed in *E. coli* Rosetta and purified under denaturing conditions using Ni-NTA columns. After separation of the concentrated protein fractions on a 10% SDS gel, rabbits were immunized with Coomassie-stained CARP1 containing gel slices according to a standard immunization protocol (Pineda, Berlin). The CARP1 antiserum was affinity-purified using His_10_-CARP1 according to the method of Olmsted. (Olmsted, 1981)

### Pull-down with cAMP agarose

1 × 10^8^ *T. brucei* cells were washed twice with PBS and lysed in lysis buffer (50 mM Tris-HCl pH 7.5, 150 mM NaCl, 2 mM EGTA, 0.2% NP-40, Roche Complete protease inhibitor) for 20 min at 4 °C. For *E. coli* lysates, 12.5 mL of a logarithmically growing culture was lysed in lysis buffer (50 mM Tris-HCl pH 7.5, 150 mM NaCl, 1% Triton X-100, 570 µM PMSF) for 10 min at 4 °C with sonication (Bioruptor, Diagenode, 10 min on ice, 30 s on/off, high power). Soluble lysates were incubated with plain agarose beads (Biolog Bremen) for 30-60 min at 4 °C to remove proteins binding non-specifically to the bead matrix. Pull-downs were performed by incubation of the pre-cleared lysates with 30-40 µL 2-AHA- or 8-AHA-agarose (Biolog Bremen Cat. No. A054, A028) beads slurry for 30 min – 2 h at 4 °C, followed by four washes with lysis buffer. Bound proteins were eluted by boiling (5 min 95 °C) with 20 µL 2× Laemmli sample buffer.

### Dose-response cell viability assay

CpdA sensitivity of the inducible CARP knock down cell lines was assessed using the Alamar blue (resazurin) cell viability assay exactly as described previously (Gould *et al*., 2013).

### Western blot

Western blot analysis was performed as previously described (Salmon *et al*., 2012a) with modifications for detection of CARP1. Briefly, lysates of 3-5 x10^6^ trypanosomes were separated on 10% polyacrylamide gels, transferred to a PVDF membrane via semi-dry blotting and blocked with Kem-En-Tec synthetic blocking buffer for 1 h at room temperature. The blots were incubated with primary antibodies (rabbit anti-CARP1, 1:1000; mouse anti-PFR-A/C ((Kohl *et al*., 1999), 1:2000) overnight at 4 °C, followed by secondary antibody (IRDye680LT anti-rabbit and IRDye800CW anti-mouse, LI-COR, both 1:5000) detection for 1.5 h at room temperature. For detection of His-tagged proteins expressed in *E. coli*, proteins were blotted onto PVDF membranes and blocked with 5% milk for 1 h at room temperature, followed by incubation with mouse anti-His (1:1000, BioRad MCA1396GA) overnight at 4 °C and secondary antibody detection with IRDye800CW anti-mouse (1:5000, LI-COR).

### Bioinformatics

CARP sequences were retrieved from TriTrypDB release 58 (https://tritrypdb.org/tritrypdb/app), information on orthology was obtained from orthoMCL release 6.11 (https://orthomcl.org/orthomcl/app), and domain predictions were done with Superfamily 2 (https://supfam.mrc-lmb.cam.ac.uk/SUPERFAMILY/), Pfam version 35.0 (http://pfam.xfam.org/), and/or SMART (http://smart.embl-heidelberg.de/). AlphaFold2 structure predictions for CARP1 and CARP11 were retrieved from http://wheelerlab.net/alphafold/ and cyclic nucleotide binding domains predicted by Superfamily 2 (https://supfam.mrc-lmb.cam.ac.uk/SUPERFAMILY/) were superimposed with crystal structures of the known cAMP binding proteins *E. coli* CRP (PDB 4N9H) and *Bos taurus* PKARIa (PDB 1RGS) using Pymol (*The PyMOL Molecular Graphics System, version 2.5* Schrödinger, LLC). Predicted CNBD phosphate binding cassette sequences of CARP1 and CARP11 were aligned with those of *E. coli* CRP, *Mus musculus* Epac4 and *Bos taurus* PKARIa using CLC Main Workbench version 21.0.5 (Qiagen).

## Acknowledgements

We thank Frank Schwede (Biolog, Bremen) for cAMP agarose beads and advice and the Tryptag.org team for a public resource that was invaluable for interpretation of the CARPs’ phenotype. Work in Munich was funded by Marie Curie PIEF-GA-2012-626034 “TrypCARPinteractors” (M.K.G) and BioNa young scientists award of the Faculty of Biology LMU (S.B.). M.A.A. was funded by a studentship from the Ministry of Higher Education of Saudi Arabia. J.C.M. was supported by an FP7 grant from the European Commission (project reference 602666). D.H. is supported by a Wellcome Trust Investigator Award [217105/Z/19/Z]. The authors are grateful to Graham Hamilton of the Glasgow Polyomics facility and Jon Wilkes of the Wellcome Centre for Molecular Parasitology of the University of Glasgow for expert assistance with the RIT-sequencing and the mapping of the reads to the genome, respectively.

## Author contributions

M.B., D.H. and H.P.d.K. designed and supervised research; S.B., M.K.G., E.P., R.O., A.E.B., M.A.A., J.C.M. performed research; S.B., M.K.G., H.P.d.K., M.B. and D.H. analyzed data; H.P.d.K., M.B. and S.B. wrote the paper.

## Data availability statement

The RIT-seq data was submitted to the European Nucleotide Archive (https://www.ebi.ac.uk/ena/browser/home), with submission number PRJEB60531.

## Additional Information / Competing interests statement

The authors declare no competing interests.

## Supplementary figure legends

**Supplementary Figure 1.**
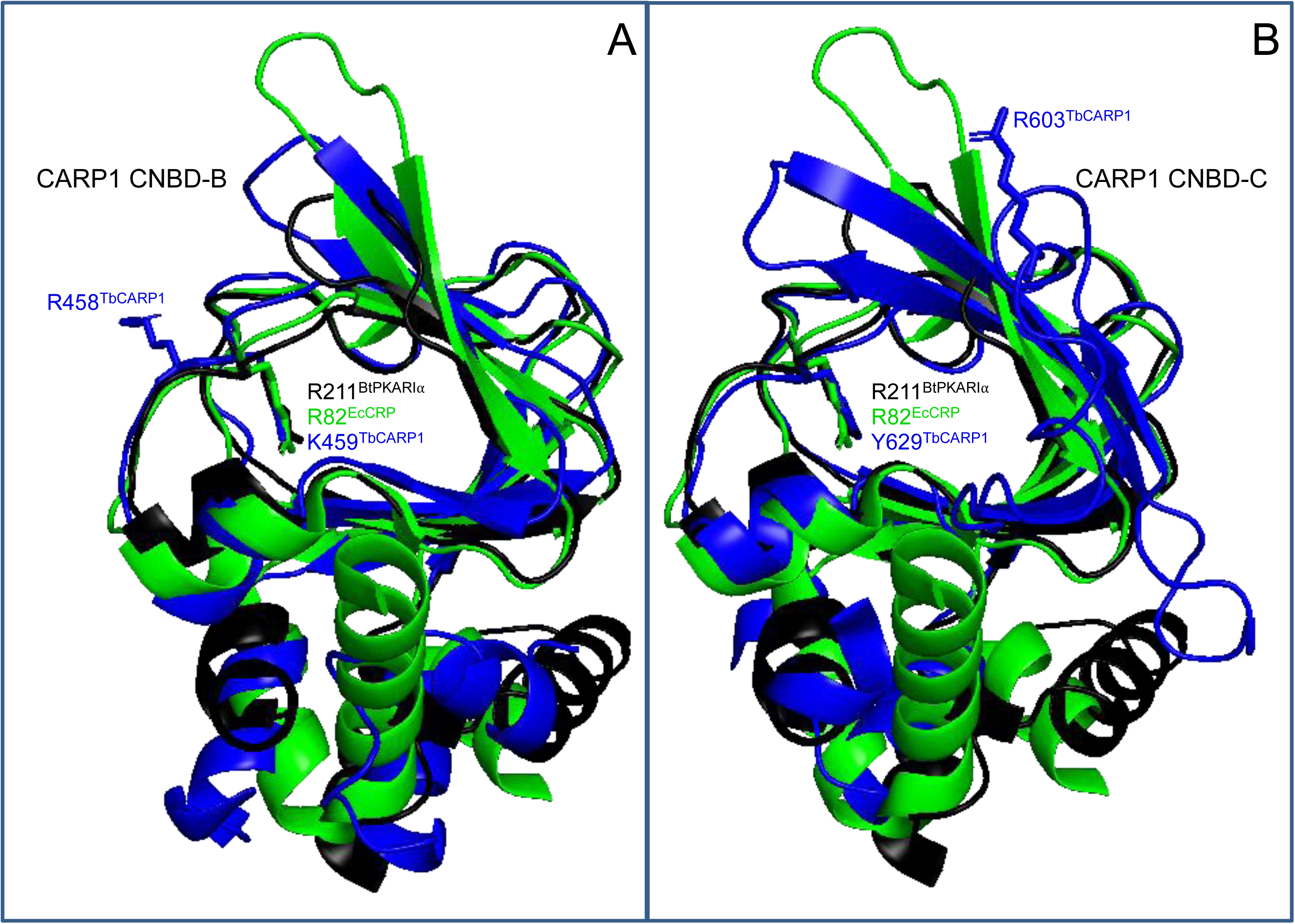
Structural alignment of CARP1 CNBD-B (A) and CNBD-C (B) with BtPKARIα_CNBD-A (PDB 1RGS, black; conserved R211 sidechain shown) and EcoliCRP_CNBD (PDB 4N9H, green; conserved R82 sidechain shown). The CARP1 CNBDs are coloured in blue. In addition to the sidechain of the Arg that is the best alignment fit in Fig. 5A, the residues that structurally align with R211^BtPKARI⍺^ and R82^EcCRP^ are displayed. The CARP1 structure predictions were retrieved from http://wheelerlab.net/alphafold/ and superimposed with Pymol version 2.5.4.

**Supplementary Figure 2.**
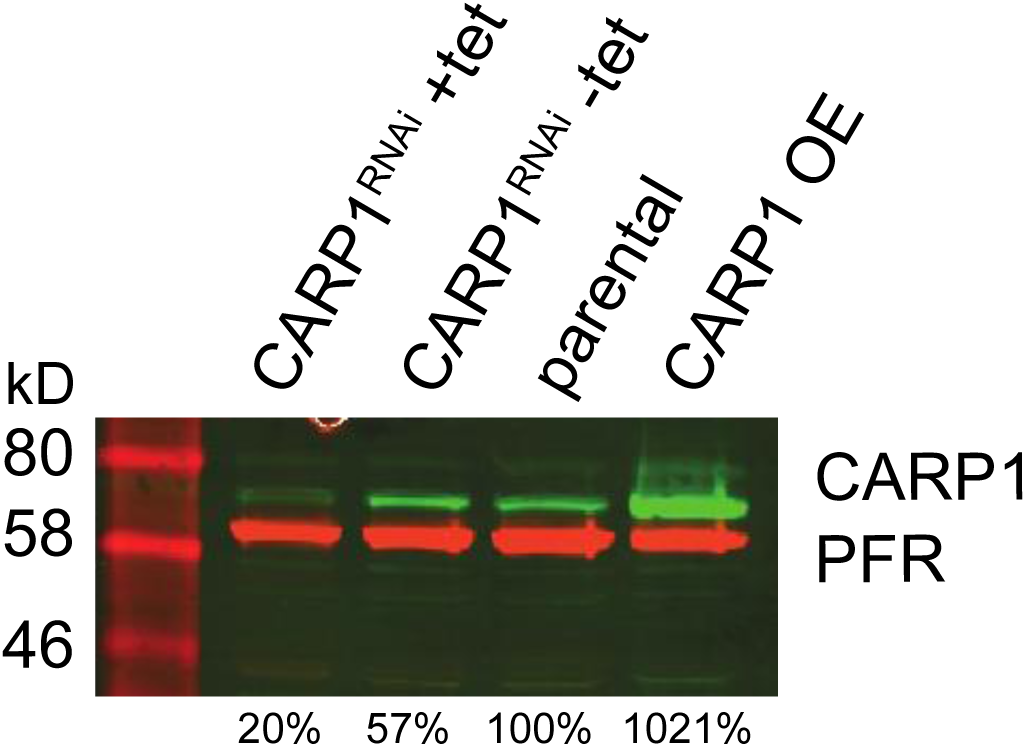
Validation and specificity of the affinity-purified rabbit anti-CARP1 antibody by western blot analysis. CARP1 expression was analyzed in trypanosome cell lines with tetracycline-inducible *CARP1* RNAi (+tet/-tet) or CARP1 overexpression (OE) compared to parental 13-90 cells. CARP1 signals (green) were normalized to the PFR-A/C loading control (red) and the normalized signal in the parental cells was set to 100%; quantification is given below the western blot.

**Supplementary Figure 3.**
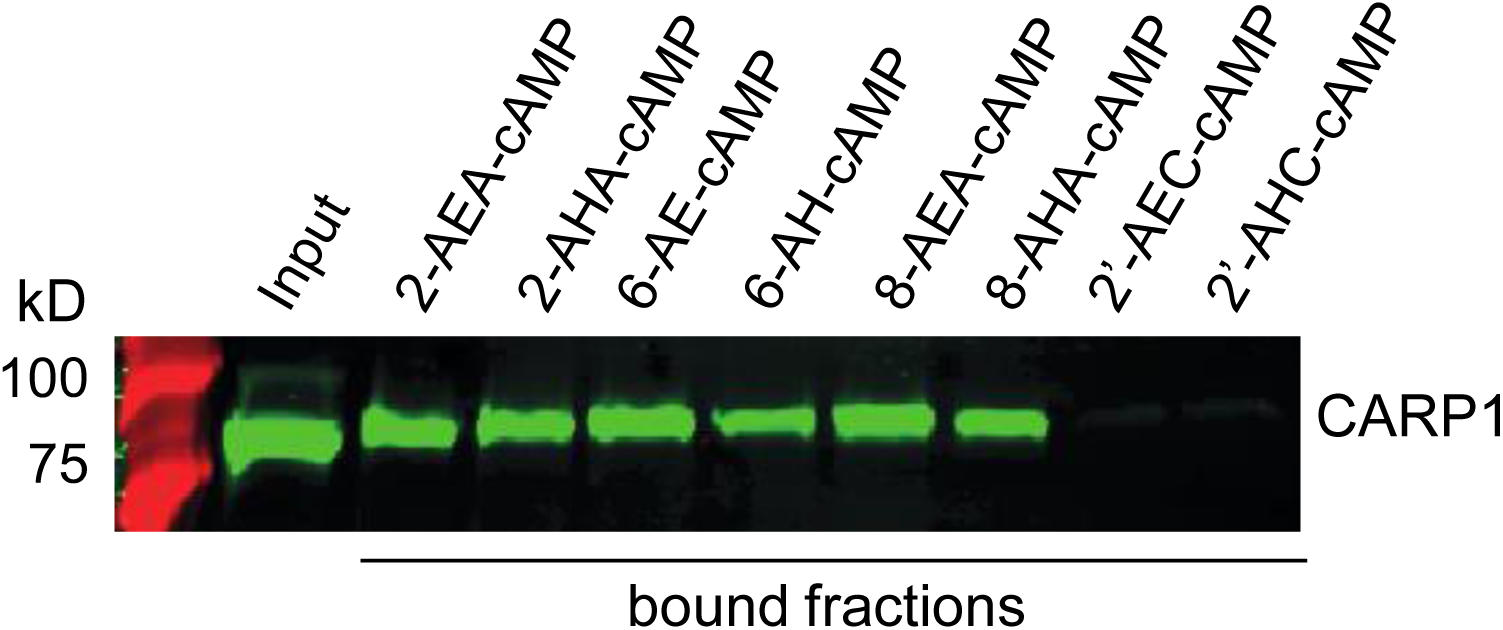
CARP1 pull-down from *T. brucei* cells overexpressing CARP1 with cAMP agarose beads with different linkers as indicated. The western blot shows the input fraction and the pulled down material (bound fractions) and was probed with rabbit anti-CARP1 (green).

